# Complete metamorphosis and microbiota turnover in insects

**DOI:** 10.1101/2021.06.15.448481

**Authors:** Chrisitin Manthey, Paul Johnston, Jens Rolff

## Abstract

The insects constitute the majority of animal diversity. Most insects are holometabolous: during complete metamorphosis their bodies are radically re-organized. This re-organization poses a significant challenge to the gut microbiota, as the gut is replaced during pupation, a process that does not occur in hemimetabolous insects. In holometabolous hosts, it offers the opportunity to decouple the gut microbiota between the larval and adult life stages resulting in high beta diversity whilst limiting alpha diversity. Here we studied 18 different herbivorous insect species from 5 orders of holometabolous and 3 orders of hemimetabolous insects. Comparing larval and adult specimens, we find a much higher beta-diversity and hence microbiota turnover in holometabolous insects compared to hemimetabolous insects. Alpha diversity did not differ between holo-and hemimetabolous insects nor between developmental stages within these groups. Our results support the idea that pupation offers the opportunity to change the gut microbiota and hence facilitates ecological niche shifts. This effect of niche shift facilitation could explain a selective advantage of the evolution of complete metamorphosis, which is a defining trait of the most speciose insect taxon, the holometabola.

## Introduction

Insects are the most diverse animal taxon on earth (1, 2) and collectively comprise 50 to 70% of all living species (2, 3). More than 80% of all described insect species are homometabolous— they undergo complete metamorphosis (3) that includes a pupal stage intercalated between the larva and the adult. In the pupa, the insect body is radically remodeled. All larval organs, including the gut, are broken down and reconstructed, resulting in distinct and specialized larval and adult life stages (4–7). Complete metamorphosis is considered a key trait explaining insect diversity (8, 9, but see 10) and only evolved once, the holometabola are a monophyletic group (11). The diversification of the speciose orders of the holometabolous insects coincides with the diversification of the land plants (10, 11).

Exactly why this radical re-organization of the insect body has produced such an astounding radiation is not known (6). One possible explanation could be that intercalating the pupal stage decouples growth and differentiation (6, 12), allowing for efficient and competitive exploitation of ephemeral resources. Another, not mutually exclusive explanation is, that larvae and adults can occupy distinct niches (13), possibly accompanied by a change in the gut microbiota.

One of the major internal reconstructions during the pupal stage includes the replacement of the gut epithelium. From the perspective of the microbes in the gut, the epithelial replacement constitutes a dramatic habitat change. The gut microbiota, that can provide nutrition, defense and other services to the insect host, changes in density and community structure, including the elimination of particular microbes during pupation (13, 14). These changes may result from a combination of factors including the drastic anatomical and physiological transformations in the replacement gut, host immune effector induction in the metamorphic gut, bacterial competition for continued occupancy of the pupal gut, and ontogenetic habitat and diet shifts of the host. In the lepidopteran *Galleria mellonella*, the absolute abundance of the microbiota can be reduced by several orders of magnitude during metamorphosis, including the elimination of pathogenic bacteria that would otherwise persist in the adult host (14). In *Galleria*, the host immune system and the symbionts interact with the microbial community in the gut through complete metamorphosis. Observations in other taxa such Lepidoptera (6 and refs therein) as well as in some Coleoptera (15) are consistent with partial host control of the microbiota. The replacement of the gut epithelium potentially offers the insect a unique opportunity to siginficantly alter the gut microbiota, allowing an insect to acquire life stage-specific microbes (13, 14). Such gut microbiota changes during the pupal stage increase the opportunity of niche shifts (16).

Some studies have investigated changes in the gut microbiota at different stages of host development. For example, the hemimetabolous insect *Pyrrhocoris apterus* (17) hosts a very stable mid-gut community composition with six predominant taxa being consistently abundant throughout development. By contrast in a holometabolous insect, the hymenopteran *Bombus pascuorum,* Parmentier et al. (18) reported different gut microbial communities within larval and adult specimens of a wild nest. The typical core gut bacteria in the adults were absent in the larvae. Hammer, McMillan, & Fierer (19) also found distinct gut microbiota communities in the leaf-chewing larvae and nectar- and pollen-feeding adults in the lepidopteran *Heliconius erato.* Studies of the dipteran *Musca domestica* (20) and the coleopteran *Phalacrognathus muelleri* (21) have found a similar pattern.

Here, we investigated whether gut microbiota changes during the adult moult, which includes pupation in the holometabolous insects, differ between hemi- and holometabolous insects. Because of the re-organization of the gut in the pupal stage we expect (a) a significant change in bacterial species composition resulting in much greater betadiversity in holometabolous than in hemimetabolous insects. The diversity of gut microbes can be strongly reduced during pupation (14, 19). (b) We therefore speculated that greater alpha diversity would be observed in the gut microbiota of hemimetabolous insects, given the lack of gut epithelial replacement. Also, as the diversity of the gut microbiota scales positively with size across species Sherill-Mix et al. (22), it is possible that alpha diversity is higher adult than larval insects, especially in hemimetabolous species.

To address these questions, we sampled 18 different species from seven major insect orders across the life stages. To reduce geographic variance, we collected all insect species in Central and Northern Europe. To further reduce variance, we sampled only herbivorous insects from terrestrial habitats and excluded social insects. Additionally, we collected a sub-sample of those species from laboratory-reared colonies, consisting of five species from four different insect orders (figure 1) as the gut microbiota may differ between specimens from the field and laboratory (23, 24).

**Figure 1:**
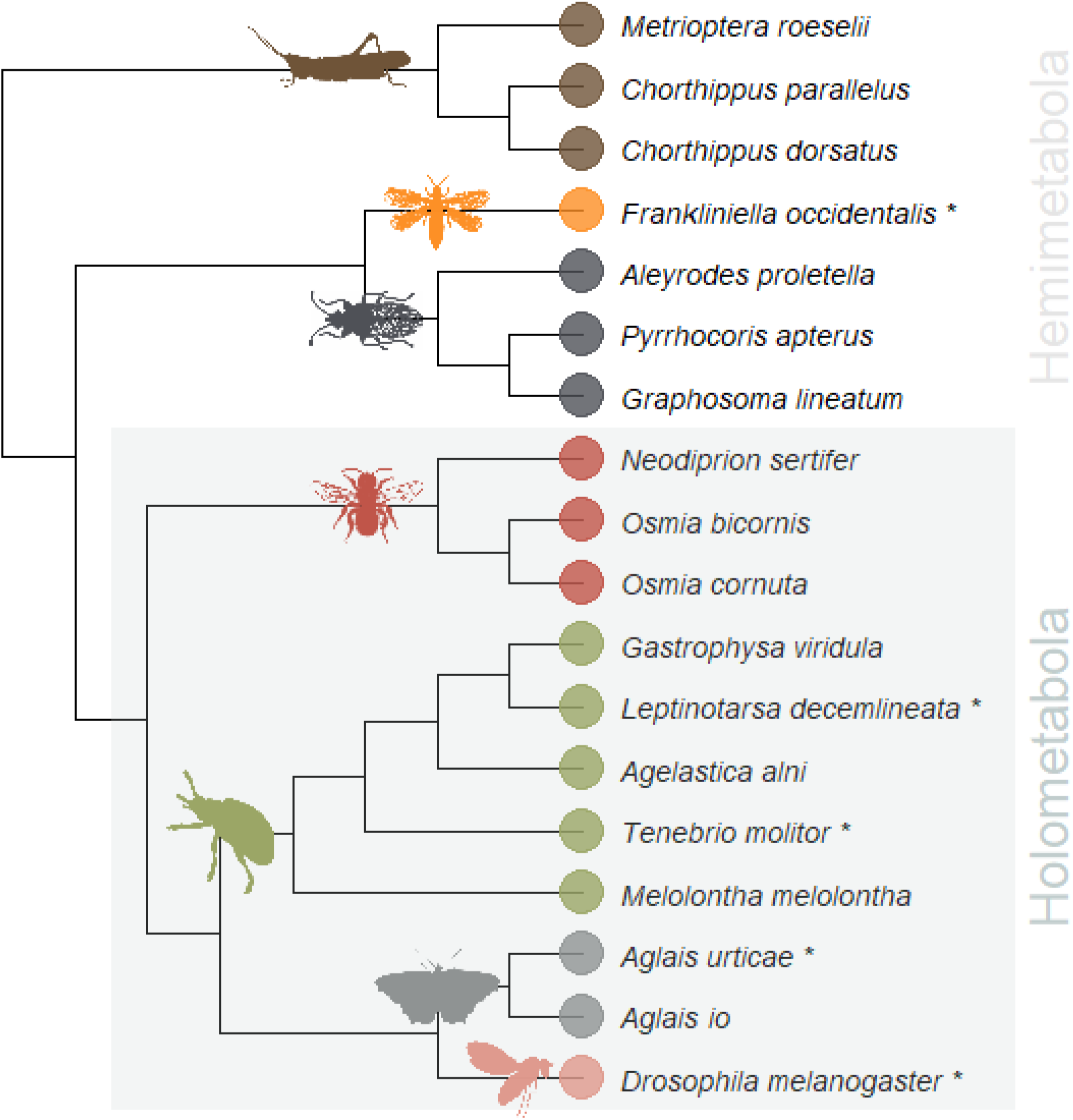
Phylogenetic tree of the 18 field-collected species. The five species marked with a star were additionally sampled from laboratory-reared colonies. The 11 highlighted species represent the monophyletic group of the holometabolous insects. The other seven species are hemimetabolous.

## Results

We obtained 18 insect species covering seven major orders, including three hemimetabolous (Orthoptera, Thysanoptera, Hemiptera) and four holometabolous insect orders (Hymenoptera, Coleoptera, Lepidoptera and Diptera). We sampled larval and adult life stages for all of them. We collected a sub-sample of those 18 species from laboratory-reared colonies that covered five insect species from four different orders, including one hemimetabolous (Thysanoptera) and four holometabolous insect orders (Coleoptera, Lepidoptera and Diptera). (Figure 1)

Using 16S rRNA sequencing, we determined the gut microbial compositions per life stage and species and plotted the data. All relative abundance plots can be found in the supplementary material. The data from *Pyrrhocoris apterus* and *Melolontha melolontha*, which originated from two different locations, were pooled as population did not affect alpha diversity (see table 3) nor beta diversity (table 2).

### Microbiota turnover at the larval-adult transition

Microbial beta diversity of holometabolous insects was significantly greater than hemimetabolous insects when comparing larval and adult life stages of each species (Two-sample t-test, t = −3.292, df = 14, p = 0.005347, figure 2). With the exception of *Leptinotarsa decemlineata*, all field-collected holometabolous insect species showed significant differences in beta diversity between larval and adult life stages (table 2).Within the hemimetabolous species collected in the field, two species differed significantly in beta diversity: *Chorthippus dorsatus* and *Pyrrhocoris apterus*. The other five Hemimetabola did not differ in beta diversity between life stages. This pattern was consistent in the subset of five laboratory-reared species. To display the differences in beta diversity between life stages for each species, we used Principal Coordinate Analysis (PCoA) ordination (see supplement, figures S1 – S18, PCoA plots per species).

**Figure 2:**
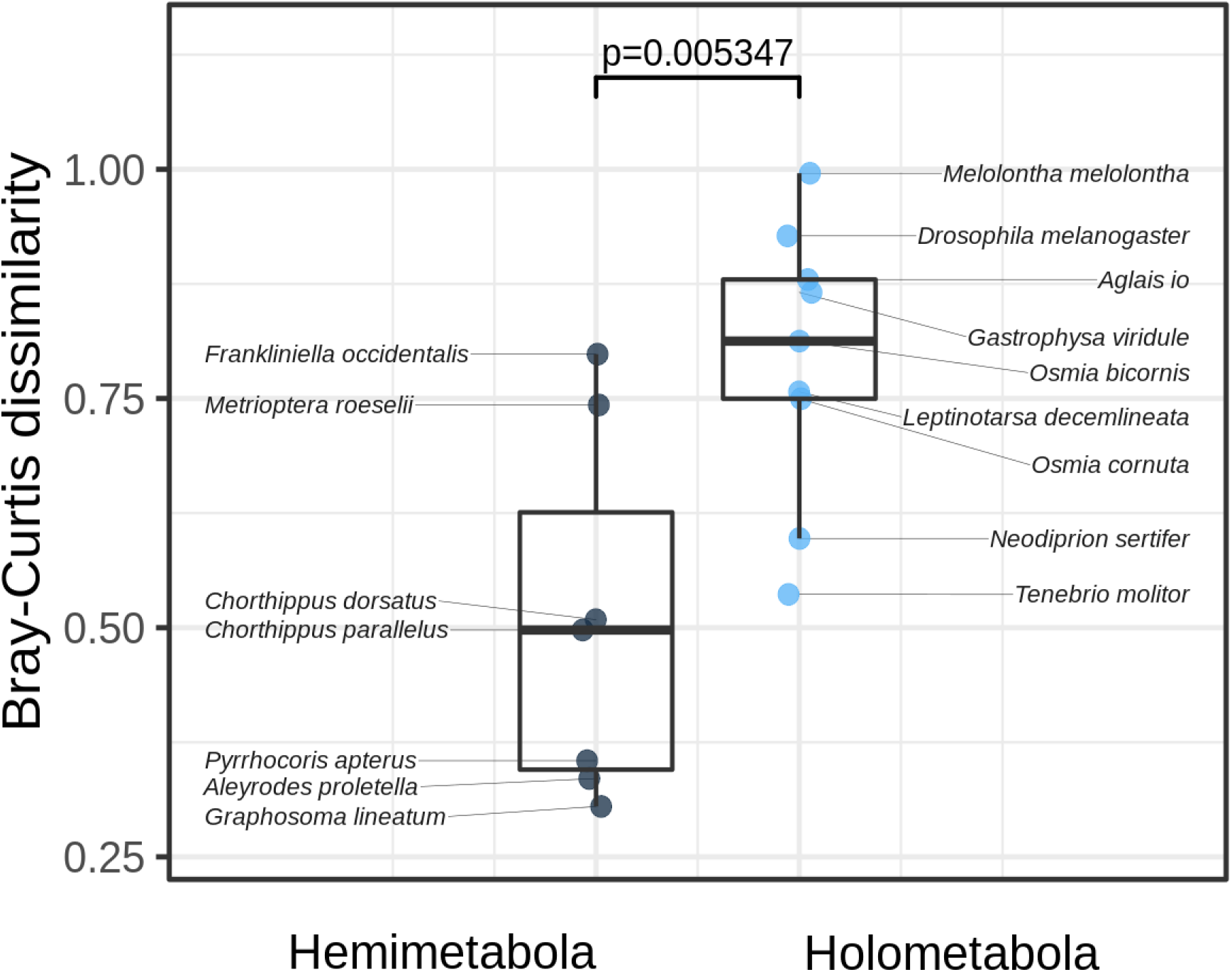
The average beta diversity (Bray-Curtis dissimilarity) in larval and adult gut communities among hemi-versus holometabolous insects (Two-sample t-test, t = −3.292, df = 14, p = 0.005347). Each point represents the beta diversity between life stages of a particular insect.

### Alpha Diversity

We calculated Shannon diversity indices per life stage and species and calculated the difference in alpha diversities between life stages per species. Alpha diversity differences between larval and adult life stages did not differ between holo- and hemimetabolous insects (Two-sample t-test, t = −1.1025, df = 14, p = 0.2889, figure 3). Because Shannon diversity does not allow to control for between-sample variation and sample sizes, we also calculated Hedges’ g effect size. Hedges’ g did not differ between holo- and hemimetabolous insects (Two-sample t-test, t = −1.3322, df = 14, p = 0.2041, figure 4). The microbial alpha diversity was also not different between holo-versus hemimetabolous larvae (Two-sample t-test, t = 0.38317, df = 14, p = 0.7073) or adults (Two-sample t-test, t = −0.18815, df = 14, p = 0.8535), respectively (figure 5). Five species did differ significantly in alpha diversity between life stages within the holometabolous insects and showed large effect sizes. Three species showed small and one species negligible effect sizes. Within the hemimetabolous insects, one species differed significantly in alpha diversity between life stages and this and a second species showed large effect sizes. The other Hemimetabola showed negligible (one species), small (three species) and medium effect sizes (one species). See table 4 for more details.

**Figure 3:**
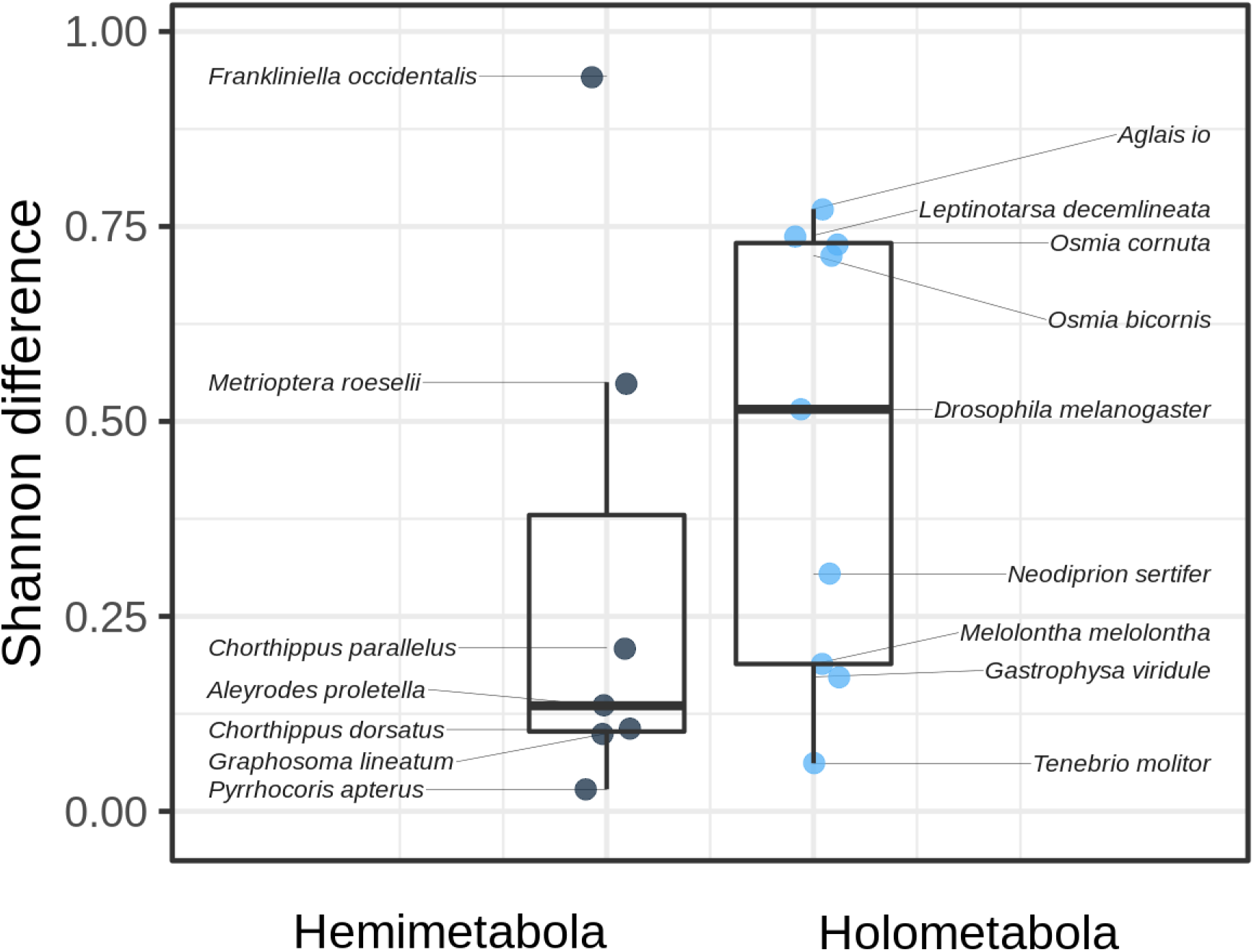
Average alpha diversity among hemi- and holometabolous insects (Two-sample t-test, t = −1.1025, df = 14, p = 0.2889). Each point represents the difference in alpha diversity, measured as Shannon difference, between larval and adult bacterial communities of a particular insect.

**Figure 4:**
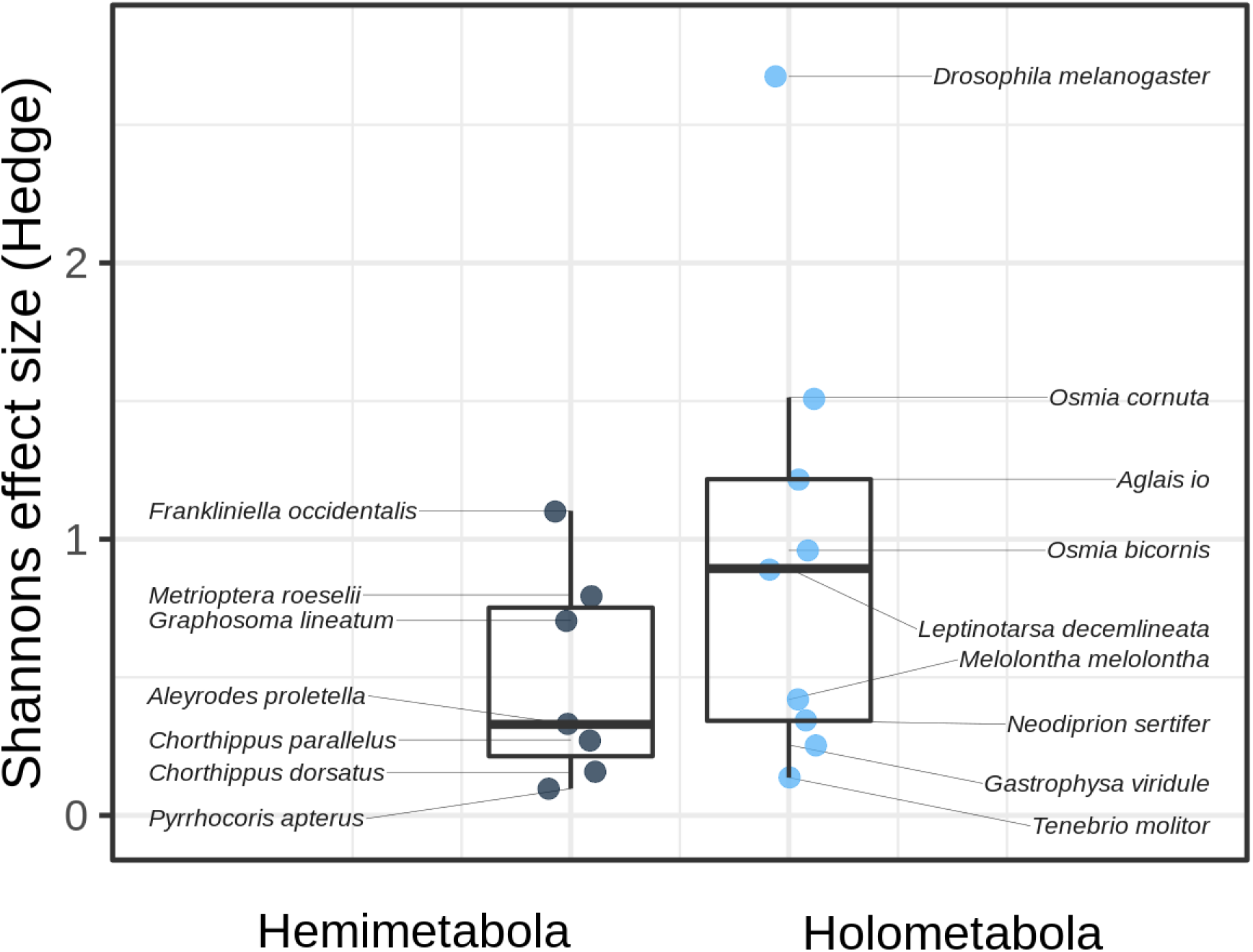
Average Shannons effect size between larval and adult gut communities among hemi- and holometabolous insects (Two-sample t-test, t = −1.3322, df = 14, p = 0.2041). Each point represents Shannons effect size, measured as Hedges’ g, between life stages of a particular insect.

**Figure 5:**
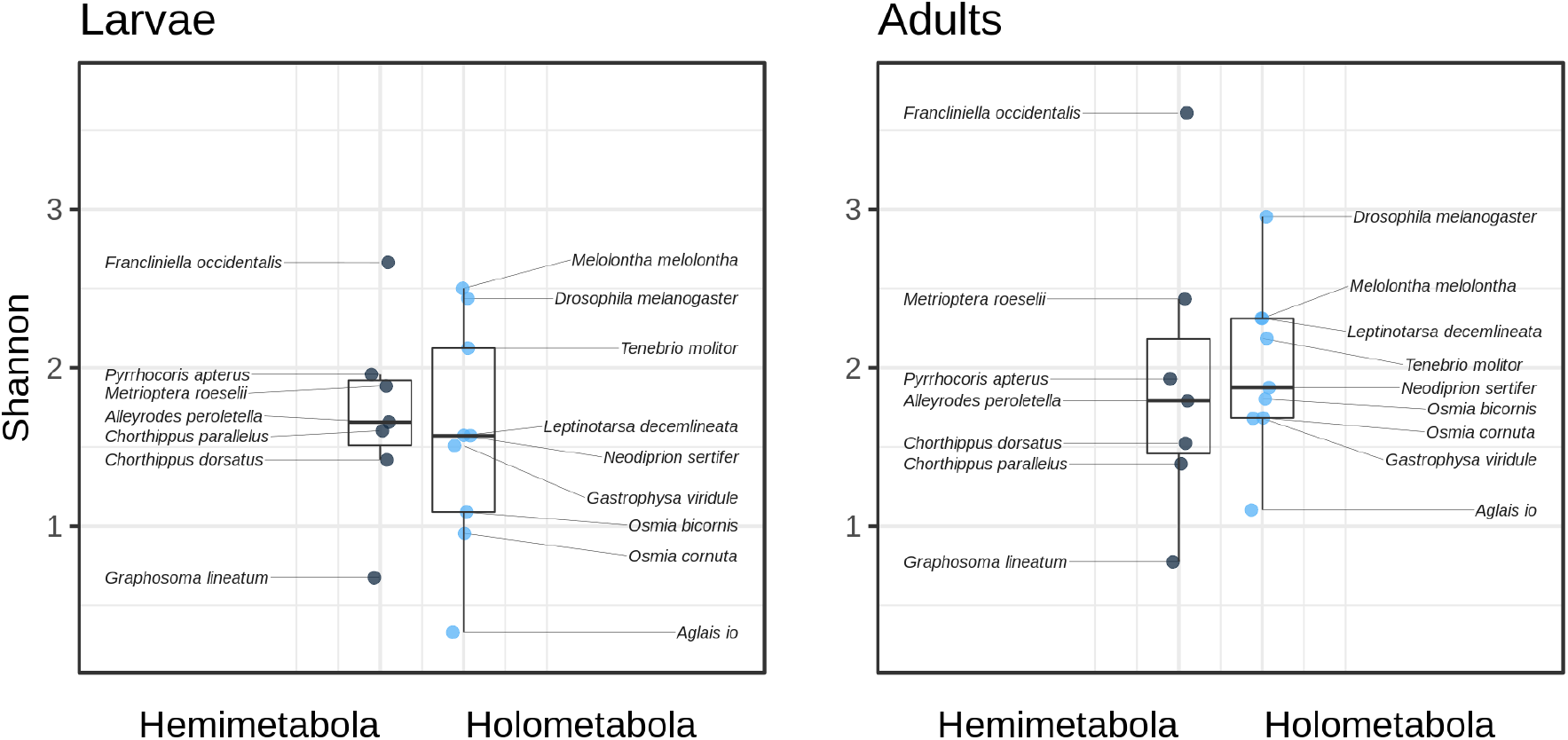
Average larval alpha diversity between hemi- and holometabolous insects (Two-sample t-test, t = 0.38317, df = 14, p = 0.7073, left), and of adults (Two-sample t-test, t = −0.18815, df = 14, p = 0.8535, right). Each point represents the alpha diversity, measured as Shannon difference, of Larvae (left) or Adults (right).

## Discussion

We investigated beta and alpha diversity throughout development, comparing 18 insect species from four holo- and three hemimetabolous insect orders. We find a clear pattern: holometabolous insects show a strong microbial turnover between larvae and adults, while this is not found in hemimetabolous insects.

Almost all examined holometabolous insect species showed significantly different gut microbial communities between larval and adult specimens as reflected by the differences in beta diversity. The overall pattern we report is well supported: hemimetabolous insects do not show changes in beta diversity during development. The remaining variation in beta diversity within the holometabolous insects may be partly explained by different ecologies which warrants further investigation. Interestingly, the highest beta diversity in hemimetabolous species was recorded in thrips. They have evolved a neometabolous life-style with two partly quiescent stages between the larva and the adult (7). The stages are also called pupae but their development does not entail the dramatic change in morphology as in holometabolous insects (7).

Microbiota turnover seems to be a general pattern within holometabolous, but not hemimetabolous insects, independent of the insects' field and laboratory origin. The microbial composition changes are presumably driven by the intercalated pupal stage in holometabolous insects, which allows a radical remodeling of the hosts' gut, but often is also accompanied by different diet choices of larvae and adults. Prior to pupation a cessation of feeding and purging of the gut contents takes place (14). After that, immune effectors such as lysozyme and AMPs are secreted into the gut (14) followed by anatomical and physiological changes resulting in the replacement of the gut. Competition of the remaining bacteria with possible new colonizers of the adult gut than shapes the adult microbiota. The role of the host immune system is illustrated by a study in *Galleria mellonella*, a species where the stage of gut replacement can be precisely determined *in vivo* (26). This study revealed that pupal gut delamination coincides with peak immune gene expression in the gut. This is consistent with other observations in other holometabolous insects with lower temporal resolution (27–29). In contrast, no such effect was observed for the hemimetabolous *Gryllus bimaculatus (26)*. The massive up-regulation of immune genes during gut renewal is therefore a candidate mechanism contributing to the patterns reported in our study. In principal, the insect host can establish a completely new and distinct adult gut microbiome by this reduction of the gut microbiota and a subsequent change in diet of the emerging adult. A different diet will expose the insects to new microbes that can colonize the gut and potentially also facilitate better digestion of the new diet.

Alpha diversity did not display a pattern related to holo- vs. hemimetabolous development. Complete metamorphosis results in a reduction of the microbial absolute abundance by orders of magnitude (14 and refs therein). In the light of Hammer *et al*. (19) this could be explained by a recovery of the microbiota upon adult feeding. Another possible explanation for the lack of change in alpha diversity are priority effects, the order and timing of arrivals determining the establishment of a new microbial community. Therefore, the gut microbiota is shaped by a niche modification, in which early arriving species change the types of niches available within the local sites (30).

The high turnover though poses the risk of losing beneficial microbes which would result in a cost to both, host and symbiont. Hammer & Moran (13) suggest that holometabolous insects may be less likely to evolve strictly vertically transmitted symbioses than hemimetabolous insects. To overcome this hurdle a number of strategies have evolved to ensure transmission of obligate symbiont between life stages in holometabolous insects. Stoll, et al (31) showed vertical transmission of microbes via bacteriocytes in ant species. The relative number of bacteria-filled bacteriocytes increased strongly during complete metamorphosis. Maire et al. (32) also showed a transmission of microbes via bacteriocytes in weevils by maintaining and relocating bacteriocytes during gut renewal in the pupa. Other specialized structures to transmit symbionts in insects are antennal glands (33) or crypts (34).

It has been suggested that microbiota turnover would allow insects to occupy different niches throughout development (13), which most likely contributed to the success of holometabolous insects. Our data are consistent with this hypothesis, the clearance of the gut provides the opportunity for a microbiota turnover, an effect not observed in hemimetabolous insects. It seems possible that this observation is directly related to the decoupling hypothesis, which proposes that growth is confined to the larval stage, while most differentiation occurs in the pupa (6, 12). A facilitation of niche shifts by changes in the gut microbiota, if confirmed by experimental studies, could be considered as an important driver of the evolution of complete metamorphosis. How this relates to the diversification of the main insects orders in relation to the diversification of land plants remains to be seen.

## Materials and Methods

### Insect Sampling and Preparation

Larval and adult specimens of 18 insect species from seven different insect orders, including Orthoptera, Thysanoptera, Hemiptera, Hymenoptera, Coleoptera, Lepidoptera and Diptera, were sampled in Central and Northern Europe between April and October 2018 (see figure 1 and table 1). Pupae were additionally sampled for the three hymenopteran species. In total, sixteen species across development were collected in Northern Germany, *Tenebrio molitor* in Croatia and *Neodiprion sertifer* in Finland. Additionally, a sub-sample of those species was sampled, consisting of five species from four different insect orders, which originated from laboratory-reared colonies. Figure 1 gives an overview of all insect species collected in the field and the subset of those species from laboratory-reared colonies. A total of 643 individual insects were sampled. All species were identified using common identification keys and were confirmed by specialists.

**Table 1:**
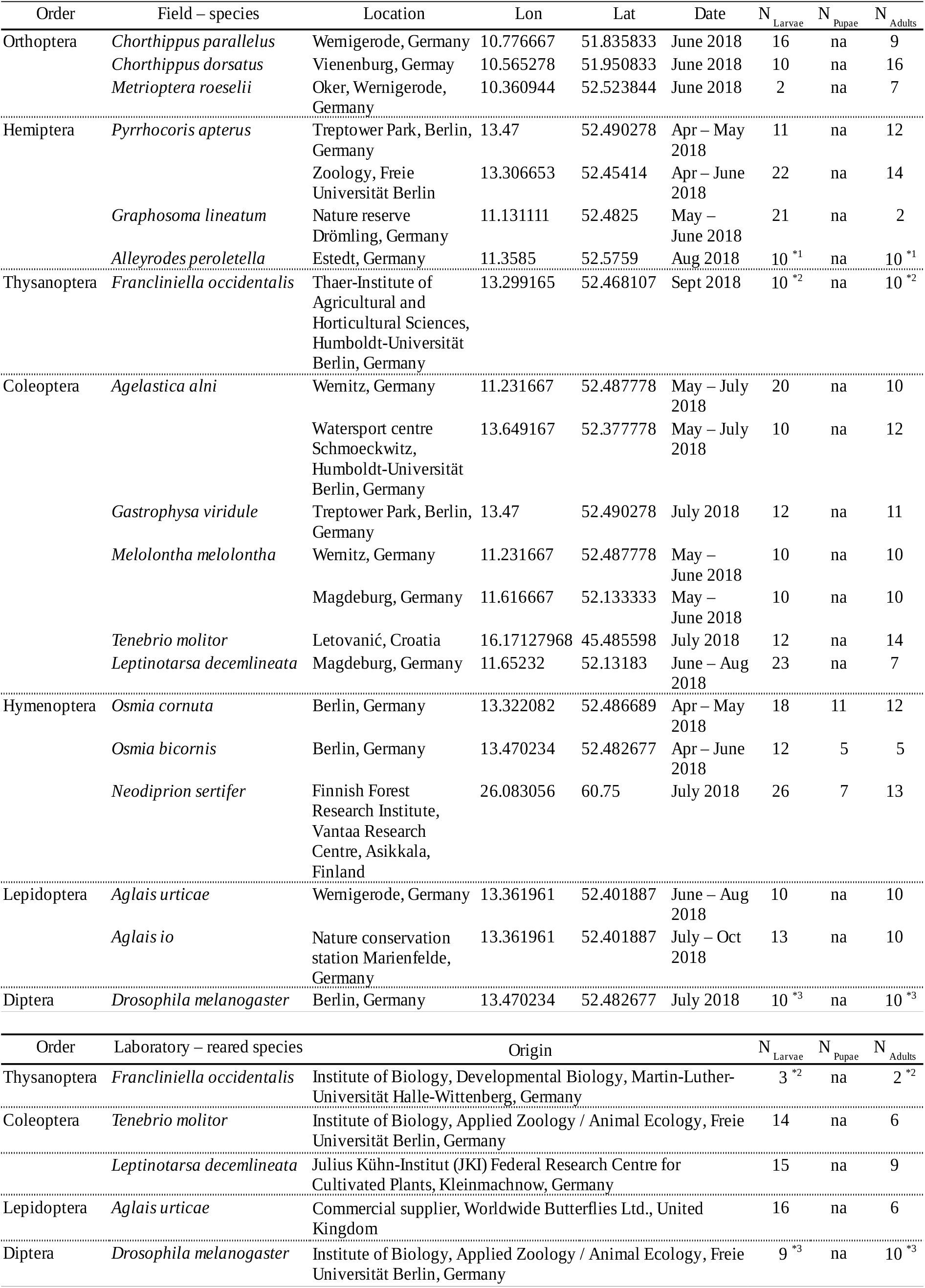
Locations from which the insects where collected. Note: ^*1^ 1 replicate equals 40 individuals, ^*2^ 1 replicate equals 10 individuals, ^*3^ 1 replicate equals 30 individuals

**Table 2:**
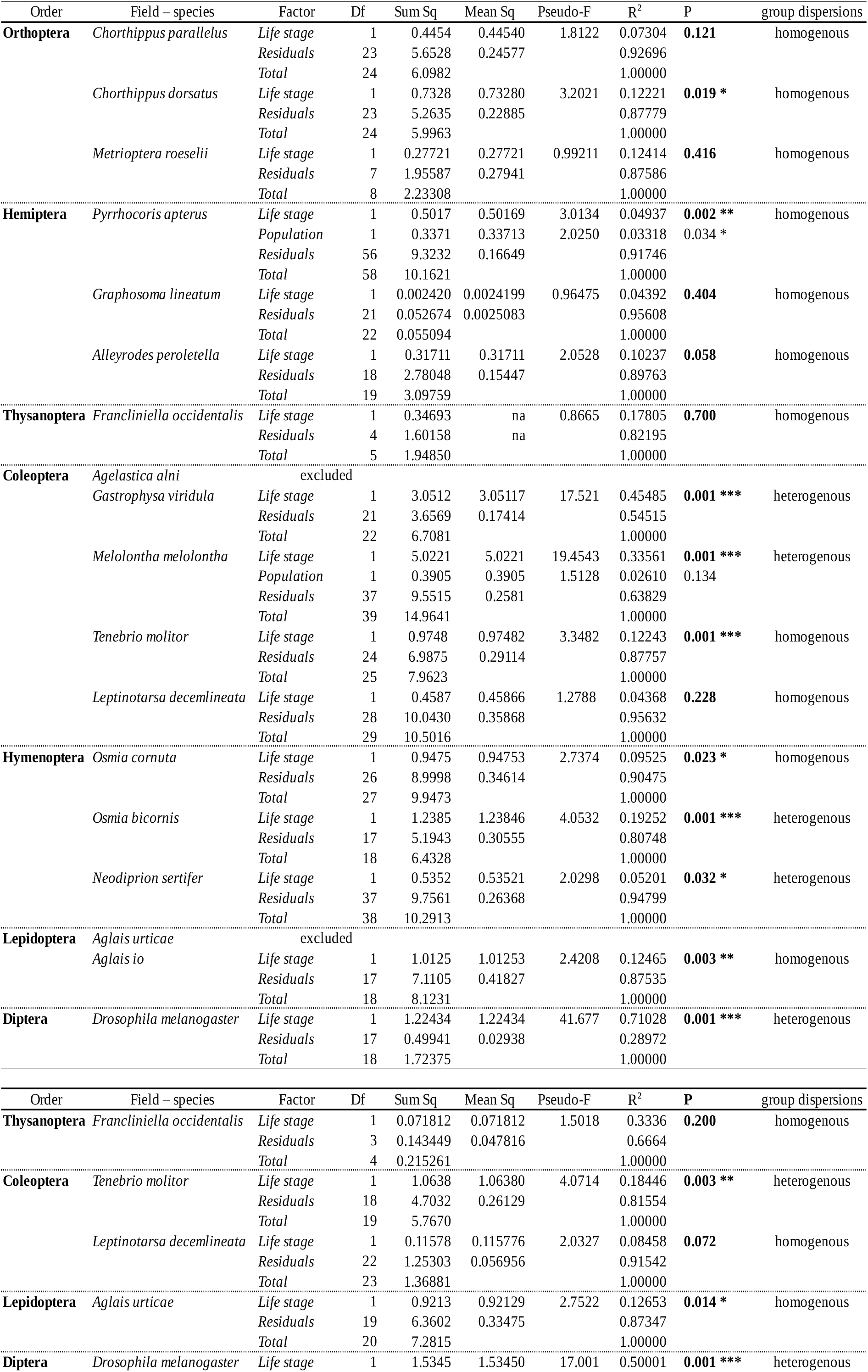
Results of perMANOVA testing differences in Beta diversity between life stages

**Table 3:** Analysis of variance table for linear model fits to compare alpha diversities between larval and adult life stages and populations per species. Shown are the species which originated from two different locations.

| Order | Field – species | Factor | Df | Sum Sq | Mean Sq | Pseudo-F | P |
| --- | --- | --- | --- | --- | --- | --- | --- |
| **Hemiptera** | *Pyrrhocoris apterus* | Life stage | 1 | 0.0116 | 0.011604 | 0.1425 | 0.7072 |
|  |  | Population | 1 | 0.1827 | 0.182706 | 2.2443 | **0.1397** |
|  |  | Residuals | 56 | 4.5589 | 0.081408 |  |  |
| **Coleoptera** | *Melolontha melolontha* | Life stage | 1 | 0.3567 | 0.35672 | 1.7861 | 0.1896 |
|  |  | Population | 1 | 0.0053 | 0.00532 | 0.0266 | **0.8713** |
|  |  | Residuals | 37 | 7.3897 | 0.19972 |  |  |

**Table 4:**
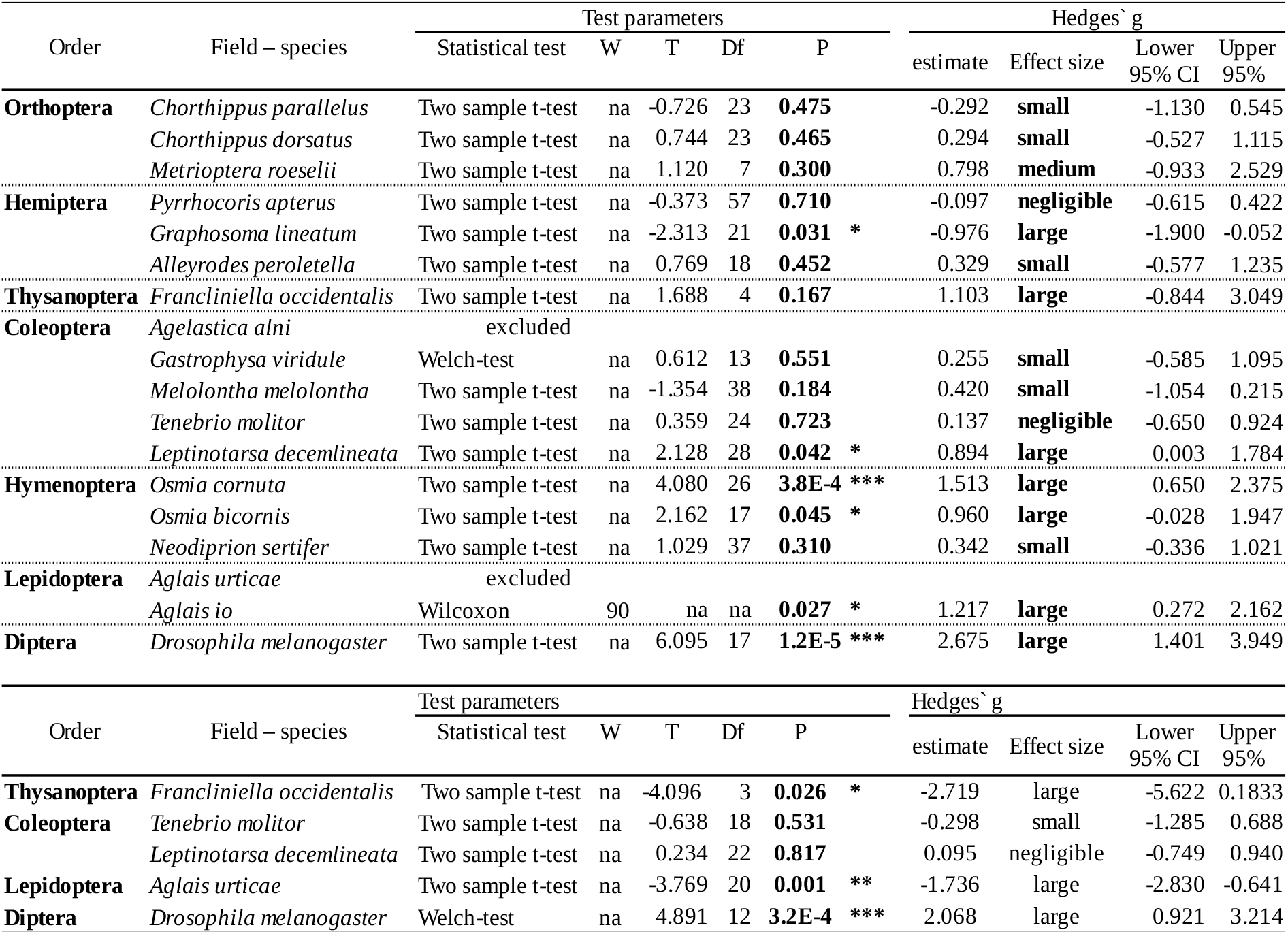
Results of t-tests used to compare larval and adult alpha diversities (Shannon). Also listed are the effect sizes (Hedges′ g).

**Table 5:** Shannon indices and evenness

| Order | Field – species | Life stage | N | Hs | SD (Hs) | E | SD (E) |
| --- | --- | --- | --- | --- | --- | --- | --- |
| **Orthoptera** | *Chorthippus parallelus* | Larva | 16 | **1.602** | 0.775 | **0.668** | 0.175 |
|  |  | Adult | 9 | **1.393** | 0.671 | **0.662** | 0.175 |
|  | *Chorthippus dorsatus* | Larva | 10 | **1.418** | 0.745 | **0.707** | 0.169 |
|  |  | Adult | 15 | **1.523** | 0.587 | **0.793** | 0.155 |
|  | *Metrioptera roeselii* | Larva | 2 | **1.884** | 0.515 | **0.744** | 0.187 |
|  |  | Adult | 7 | **2.434** | 0.627 | **0.765** | 0.075 |
| **Hemiptera** | *Pyrrhocoris apterus* | Larva | 33 | **1.957** | 0.292 | **0.782** | 0.082 |
|  |  | Adult | 26 | **1.929** | 0.284 | **0.819** | 0.059 |
|  | *Graphosoma lineatum* | Larva | 8 | **0.676** | 0.024 | **0.802** | 0.214 |
|  |  | Adult | 15 | **0.775** | 0.166 | **0.707** | 0.165 |
|  | *Alleyrodes peroletella* | Larva | 10 | **1.656** | 0.362 | **0.875** | 0.125 |
|  |  | Adult | 10 | **1.791** | 0.423 | **0.868** | 0.052 |
| **Thysanoptera** | *Francliniella occidentalis* | Larva | 3 | **2.664** | 0.892 | **0.858** | 0.057 |
|  |  | Adult | 3 | **3.607** | 0.373 | **0.891** | 0.020 |
| **Coleoptera** | *Agelastica alni* | excluded |  |  |  |  |  |
|  | *Gastrophysa viridule* | Larva | 12 | **1.508** | 0.354 | **0.813** | 0.076 |
|  |  | Adult | 11 | **1.680** | 0.870 | **0.648** | 0.250 |
|  | *Melolontha melolontha* | Larva | 20 | **2.501** | 0.476 | **0.887** | 0.040 |
|  |  | Adult | 20 | **2.312** | 0.403 | **0.778** | 0.087 |
|  | *Tenebrio molitor* | Larva | 12 | **2.123** | 0.466 | **0.784** | 0.089 |
|  |  | Adult | 14 | **2.185** | 0.409 | **0.775** | 0.099 |
|  | *Leptinotarsa decemlineata* | Larva | 23 | **1.572** | 0.761 | **0.778** | 0.150 |
|  |  | Adult | 7 | **2.311** | 0.944 | **0.768** | 0.096 |
| **Hymenoptera** | *Osmia cornuta* | Larva | 16 | **0.954** | 0.407 | **0.723** | 0.149 |
|  |  | Pupa | 8 | **1.268** | 0.611 | **0.699** | 0.122 |
|  |  | Adult | 12 | **1.683** | 0.540 | **0.843** | 0.104 |
|  | *Osmia bicornis* | Larva | 11 | **1.089** | 0.279 | **0.744** | 0.126 |
|  |  | Pupa | 5 | **1.011** | 0.335 | **0.814** | 0.172 |
|  |  | Adult | 8 | **1.802** | 1.054 | **0.790** | 0.143 |
|  | *Neodiprion sertifer* | Larva | 26 | **1.570** | 0.941 | **0.712** | 0.179 |
|  |  | Pupa | 7 | **2.123** | 0.497 | **0.748** | 0.086 |
|  |  | Adult | 13 | **1.874** | 0.701 | **0.665** | 0.131 |
| **Lepidoptera** | *Aglais urticae* | excluded |  |  |  |  |  |
|  | *Aglais io* | Larva | 13 | **0.330** | 0.463 | **0.752** | 0.390 |
|  |  | Adult | 9 | **1.102** | 0.782 | **0.551** | 0.352 |
| **Diptera** | *Drosophila melanogaster* | Larva | 10 | **2.437** | 0.170 | **0.908** | 0.024 |
|  |  | Adult | 9 | **2.953** | 0.199 | **0.900** | 0.025 |
| Order | Field – species | Life stage | N | Hs | SD (Hs) | E | SD (E) |
| **Thysanoptera** | *Francliniella occidentalis* | Larva | 3 | **2.067** | 0.338 | **0.633** | 0.062 |
|  |  | Adult | 2 | **0.948** | 0.200 | **0.462** | 0.016 |
| **Coleoptera** | *Tenebrio molitor* | Larva | 14 | **1.377** | 0.793 | **0.641** | 0.273 |
|  |  | Adult | 6 | **1.158** | 0.363 | **0.732** | 0.164 |
|  | *Leptinotarsa decemlineata* | Larva | 15 | **1.515** | 0.295 | **0.768** | 0.077 |
|  |  | Adult | 9 | **1.545** | 0.324 | **0.751** | 0.103 |
| **Lepidoptera** | *Aglais urticae* | Larva | 16 | **1.585** | 0.609 | **0.760** | 0.220 |
|  |  | Adult | 6 | **0.541** | 0.475 | **0.883** | 0.163 |
| **Diptera** | *Drosophila melanogaster* | Larva | 9 | **0.940** | 0.212 | **0.580** | 0.118 |
|  |  | Adult | 10 | **1.758** | 0.479 | **0.704** | 0.126 |

After collection, the insects were stored individually in 50 ml centrifuge tubes (Falcon tubes) with holes for ventilation. In two very small species, *Frankliniella occidentalis* and *Aleyrodes proletella*, individuals were pooled (see table 1). These pools were kept and used as a biological replicate later. After a 24-hour starvation period, the insects were sacrificed, and preserved by freezing (−80°C), except two field-collected species which were preserved in ethanol (95%): *Frankliniella occidentalis* and *Neodiprion sertifer*. Hammer, Dickerson, & Fierer (35) compared two different storage methods, freezing and ethanol, amongst others, and found that the storage method did not affect microbiota composition assessments.

### DNA extraction

Samples of the sacrificed insects were processed on ice under sterile conditions. A biological replicate was an individual insect sample, except for samples of three small species: a replicate of *Frankliniella occidentalis* was pooled from 30 individuals, a replicate of *Aleyrodes proletella* from 40 and a replicate of *Drosophila melanogaster* from 10 individuals. The exact number of biological replicates per species and life stage are shown in table 1. After cutting the legs and wings off (only adults), the samples were placed in 2 ml microcentrifuge tubes (Eppendorf Safe-Lock Tubes). Then samples were bead-ground using TCBeads and C1 solution from the PowerSoil DNA Isolation Kit (Qiagen) three times for 30 sec at 30 Hz in a tissue homogenizer. The insects were not dissected before homogenization in order to process all samples under standardized methods as the Thrips and Whiteflies were too small to dissect guts. Insects were not sterilised prior to homogenisation. Hammer et al. (35) found no effect on the bacterial communities of not surface sterilised insect species (butterfly, grasshopper, bee and beetle) compared to control specimens that were surface sterilised. Samples clustered by species independent of surface sterilisation and relative abundances of bacterial genera were similar between sterilised and non-sterilised specimens. Also, surface contaminants derived from handling the specimens were extremely rare in non-sterilised and surface sterilised specimens. As it remains possible that surface sterilisation could affect internal bacterial communities, Hammer et al. (35) recommend omitting surface sterilisation from insect microbiota studies.

Total DNA was extracted from 60 μl of tissue homogenate using the PowerSoil DNA extraction kit (Qiagen) under sterile conditions. Tissue homogenates were pretreated with 10 μl Proteinase K and 500 μl Power soil bead solution at 56 °C overnight. Subsequent DNA isolation was continued as indicated in the manufacturer‘s instruction.

Negative extraction controls were included to detect and filter contamination. The negative controls consisted of mock samples, which contained no insect tissue.

### Primers and PCR amplification of the 16S rRNA gene fragment

PCR amplification of the 16S rRNA gene fragments was performed with MyTaqTM HS DNA Polymerase and the forward and reverse 515f-806r primer sequence pairs, targeting the V3 – V4 region of the 16S rRNA gene (36). The PCR reaction was conducted using 1 μl sample in a total volume of 25 μl. The PCR amplification program was as follows: 94 °C for 1 min, 95 °C for 15 sec, 50 °C for 15 sec, two cycles of 72 °C for 45 sec and 2 min, followed by a final extension step to 4 °C. A volume of 5 μl of the PCR product was run on a 1.5% agarose gel stained with Sybr Gold at 160 V for 40 min.

PCR products were purified with CleanNGS CNGS-0050 (GC biotech B.V., Leidse Schouw 2, 2408 AE Alpen aan den Rijn, Netherlands) and dual indices and Illumina sequencing adapters were attached by limited-cycle PCR amplification (initial denaturation at 95°C for 2 min followed by eight cycles of denaturation at 95°C for 20 s, annealing at 52°C for 30 s, extension at 72°C for 30 s, and a final extension cycle at 72°C for 3 min). The enzymes used were Herculase II Fusion DNA Polymerase (Agilent Technologies Sales & Services GmbH & Co. Hewlett-Packard-Str. 831, 76337 Waldbronn, Germany). PCR products were quantified with Quant-iT™ PicoGreen™ dsDNA Assay Kit (Life Technologies GmbH Thermo Fisher Scientific,Frankfurter Straße 129b, 64319 Darmstadt, Germany), measured with Optima Fluostar (BMG LABTECH GmbH, Allmendgrün 8,77799 Ortenberg, Germany).

### Sequencing of bacterial community

Amplicon libraries were sequenced for 600 cycles using an Illumina MiSeq (Illumina, San Diego, California, USA) at the Berlin Center for Genomics in Biodiversity Research (BeGenDiv). The resulting 300-bp paired end reads were analysed using a full-stack R (37) pipeline incorporating dada2 (38) and phyloseq (39). Forward reads were trimmed to 240 bp and reverse reads to 160 bp. The reads were truncated at the first instance of a quality score less than two and filtered to a maximum amount of estimated errors of two per truncated read. The remaining forward and reverse reads were dereplicated, and error rate estimates were computed. The developed error model was used to infer exact amplicon sequence variants (ASVs) from the amplicon sequencing data. The resulting denoised read pairs were merged. A sequencing table was constructed with the denoised and merged reads and chimaeras were removed. Taxonomy was assigned to the sequence table using the Ribosomal Database Project (40) training set, version 16. Contaminant taxa were identified using prevalence-based filtering from the decontam package (41). Remaining unknown sequences were identified using the Basic Local Alignment Search Tool (BLAST) (42) and taxonomy was assigned using TaxonKit (43). Further remaining unknown sequences (genus level) were renamed with higher taxonomic ranks. Eukaryota were removed, and the resulting sequence variants were agglomerated at the genus level.

### Statistical Analysis

Absolute abundances were plotted for all species and life stages using the R package microbiome (44). Rare bacterial taxa present less than 1% of all taxa per species are not shown in the figures. The sequences were agglomerated at the genus level. Shannon indices were calculated per developmental stage and species using the microbiome package (44). To adjust for differences in library sizes, Willis (45) suggests accounting for unobserved taxa instead of rarefying the data. The breakaway estimator (46) was used but didn't differ from an estimator that does not account for unobserved taxa. Therefore the simpler approach using proportions was used. The number of reads can be found in the supplementary material. The group means per species were compared with life stage as grouping factor using, according to test assumptions, a Two-sample t-test, Wilcoxon rank-sum test and Welch-test, respectively. See table 4 for more details. To meet the assumption of normally distributed data for the Two-sample t-test, the response variable was transformed before testing for group differences using logarithm transformation for *Chorthippus parallelus* and *Chorthippus dorsatus,* and via reciprocal 1/x^6 transformation for the dataset of *Graphosoma lineatum.* The effect sizes were calculated using the effsize package (47).

Bray-Curtis dissimilarity matrices were calculated comparing life stages per species. The data was normalised to proportions to control for read depth prior to ordination. The dissimilarity data was analysed by perMANOVA with life stage as a predictor variable, and a dispersion test was fitted using the vegan package (48). Principle coordinate analysis (PCoA) on Bray-Curtis dissimilarity was used to display the beta diversities per species with life stage as a grouping factor using the phyloseq package (39).

## Acknowledgments

We are grateful to Olivia Judson and Dino McMahon for comments on the manuscript and Elisa Bittermann, Sarah Sparmann and Susan Mbedi for support in the lab. This research was funded by the DFG (Deutsche Forschungsgemeinschaft).

